# Species and Genetic Diversity of Nontuberculous Mycobacteria (NTM) in Suspected Tuberculosis Cases in East Azerbaijan, Iran: A Cross-sectional Analysis

**DOI:** 10.1101/2024.06.10.598344

**Authors:** M Roshdi Maleki

**Affiliations:** Department of Microbiology, Malekan Branch, Islamic Azad University, Malekan, Iran

**Keywords:** Genetic diversity, Nontuberculous mycobacteria, Species, Suspected tuberculosis, NTM Infections, Diagnostic

## Abstract

**Introduction:** The outbreak of nontuberculous mycobacterial (NTM) infections has increased worldwide, attracting attention in routine diagnostic settings, particularly among patients with suspected tuberculosis. This study aimed to acquire knowledge of NTM infections in patients with suspected tuberculosis and to evaluate the genetic diversity of the strains.

**Methods:** In this study, 230 clinical specimens were collected from suspected tuberculosis patients. Following decontamination with N-Acetyl-L-cysteine–sodium hydroxide (NALC-NaOH), the sediments of specimens were inoculated onto Löwenstein–Jensen medium and then incubated at 30°C for 8 weeks. The samples that yielded positive cultures underwent evaluation through sequencing conserved fragments of *IS6110* and *hsp65*. For those samples that were not identified as part of the *M. tuberculosis* complex (MTC) by *IS6110* PCR, further analysis was conducted using PCR to detect fragments of the *hsp65* gene.

**Results:** Twenty-one NTM species were isolated from 230 clinical specimens (14 NTM from pulmonary specimens and 7 from extrapulmonary specimens). Among these, 12 (57.14%) were rapid-growing mycobacteria (RGM) and 9 (42.85%) were slow-growing mycobacteria (SGM). No *M. avium* complex (MAC) was identified in any of the specimens. Notably, *M. kansasii, M. gordonae*, and *M. abscessus* strains exhibited significant genetic diversity.

**Conclusions:** The prevalence of infections attributed to nontuberculous species surpasses that of tuberculosis. These findings underscore the importance of exploring NTM species in individuals suspected of having TB.

## 1. Introduction

The genus Mycobacterium contains two major human pathogens: Mycobacterium (M.) tuberculosis (MTB), the main cause of tuberculosis (TB), and *M. leprae*, the agent responsible for leprosy. Alongside these two obligate human pathogens, the genus also includes the nontuberculous mycobacteria (NTM) or Mycobacteria other than *M. tuberculosis* (MOTT) [1].

Nontuberculous mycobacteria (NTM) represent a diverse group of opportunistic pathogenic bacteria originating from the environment. Over 170 distinct species of NTM have been recognized [2]. While the majority of NTM species are saprophytic, approximately one-third of them have been linked to human illnesses [3].

In recent years, the incidence of NTM-related diseases has significantly increased [4]. The lungs are the most common site of NTM infection, but other parts of the body such as soft tissues, blood, lymph nodes, and the skin can also be affected [5]. NTM infections are usually believed to be a result of environmental exposures, including water, soil, and dust sources [6].

Nontuberculous mycobacterial pulmonary disease (NTM-PD) was initially identified as concomitant infections among a small subset of patients in tuberculosis (TB) sanatoria. Currently, NTM-PD poses a significant challenge as its incidence is on the rise, with reasons for this trend remaining unclear. The NTM species responsible for pulmonary disease exhibit regional variability, with the *M. avium-intracellulare* complex (MAC) emerging as the primary pathogen in many areas (Table 1) [7,8].

**Table 1.**
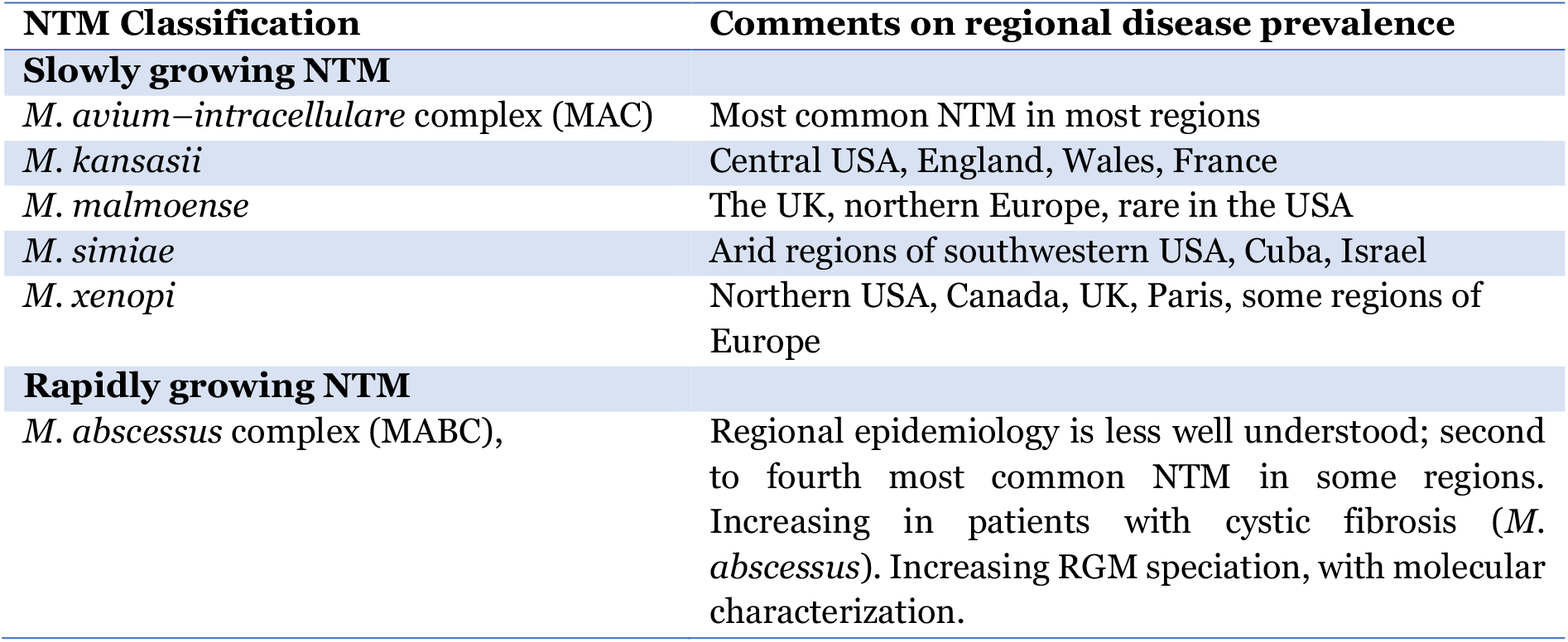
Nontuberculous mycobacteria that cause pulmonary disease: observations on regional prevalence.

| NTM Classification | Comments on regional disease prevalence |
| --- | --- |
| **Slowly growing NTM** |  |
| *M. avium–intracellulare* complex (MAC) | Most common NTM in most regions |
| *M. kansasii* | Central USA, England, Wales, France |
| *M. malmoense* | The UK, northern Europe, rare in the USA |
| *M. simiae* | Arid regions of southwestern USA, Cuba, Israel |
| *M. xenopi* | Northern USA, Canada, UK, Paris, some regions of Europe |
| **Rapidly growing NTM** |  |
| *M. abscessus* complex (MABC), | Regional epidemiology is less well understood; second to fourth most common NTM in some regions. Increasing in patients with cystic fibrosis (*M. abscessus*). Increasing RGM speciation, with molecular characterization. |

According to Runyon, NTMs can be classified into four types based on the growth rate and pigment formation (Table 2). Types I, II, and III strains are classified as slow-growing mycobacteria (SGM) because they take more than seven days to form visible colonies on a culture plate. They differ in their ability to produce pigments. The most significant species include *M. kansasii, M. malmoense, M. simiae, M. marinum*, and *M. xenopi*. Type IV strains are considered rapid-growing mycobacteria (RGM) because they take less than seven days to form visible colonies on a culture. The most clinically important species in this category are the *M. abscessus* complex (MABC), *M. chelonae*, and *M. fortuitum* [9].

**Table 2.** Classification of NTMs according to Runyon, and their reported pathogenesis in humans.

| Runyon Classification | NTM Species (e.g.) | Pathogenesis |
| --- | --- | --- |
| Photochromogens<br>Runyon type I | *M. kansasii*<br>*M. simiae* | Pulmonary infections<br>Skin infections<br>Disseminated infections |
| Scotochromogens<br>Runyon type II | *M. gordonae* | Pulmonary infections<br>Skin infections<br>Disseminated infections |
| Non-photochromogens<br>Runyon type III | MAC | Pulmonary MAC infections<br>Disseminated infections (mostly in AIDS patients)<br>MAC associated lymphadenitis (in young kids and people with normal immune systems) |
| Rapid growing<br>Runyon type IV | *M. abscessus*<br>*M. chelonae*<br>*M. fortuitum* | Skin and soft tissue infections (SSTI)<br>Pulmonary infections<br>Disseminated infections |

Several studies in different countries have reported a rise in the prevalence of pulmonary diseases over time [10,11,12,13,14]. NTM were isolated from clinical specimens of hospitalized patients, which were sent to the Mycobacteriology Research Center (MRC) at Tabriz University of Medical Sciences (TUMS) for diagnosis.

The prevalence of NTM infections has increased in patients with suspected tuberculosis. This study aimed to survey the genetic diversity of NTM species in patients with suspected tuberculosis in East Azerbaijan province, Iran. Despite the importance of NTMs, no other research has been conducted in this region, making this the first study of its kind on NTMs in northwestern Iran.

## 2. Material and methods

### 2.1. Collection of specimens and decontamination

This cross-sectional analysis involved the isolation of NTM from clinical specimens of patients with suspected pulmonary tuberculosis received between 2021 and 2022. A total of 230 clinical specimens (136 pulmonary and 94 extrapulmonary) were collected from suspected tuberculosis patients at health centers in East Azerbaijan province, Iran. The pulmonary specimens consisted of 66.2% (90/136) sputum and 33.8% (46/136) bronchoalveolar lavage fluid (BALF). The extrapulmonary specimens included 44.7% (42/94) skin samples, 36.2% (34/94) urine samples, and 19.1% (18/94) lymph node samples.

The suspected specimens underwent decontamination using the N-acetyl-L-cysteine 2% sodium hydroxide (NALC-2% NaOH) assay. Following decontamination, the sediments of the specimens were inoculated onto Löwenstein–Jensen medium and then incubated at 30°C for a duration of 8 weeks. Weekly monitoring was conducted to observe mycobacterial growth. The confirmation of suspected colonies as acid–alcohol–resistant bacilli was performed using the Ziehl-Neelsen staining technique [3,15].

### 2.2. DNA Extraction

For DNA extraction, a loop-full of mycobacterial cells was suspended in 500 μl of 1X Tris-EDTA buffer (10 mM Tris-HC1, 1 mM EDTA, pH 8.0) and was heated at 80°C for 30 min to lyse the cells. Subsequently, DNA was extracted by the cetyltrimethylammonium bromide (CTAB)/NaCl method, as described by van Soolingen et al. [16].

### 2.3. PCR for IS6110 and hsp65

The primers used (MTB1, 5′ CCTGCGAGCGTAGGCGTCGG 3′, and MTB2, 5′ CTCGTCCAGCGCCGCTTCGG 3′) amplified a 123-bp fragment of *IS6110*, which is specific for *M. tuberculosis* [17]. Samples that were not identified as *M. tuberculosis* by *IS6110* PCR were submitted to PCR for the detection of fragments of the *hsp65* gene [6, 18, 19]. In this study, the sequence analysis of the *hsp65* gene was used to identify NTMs [18]. A 441-bp fragment of the *hsp65* gene was amplified with primer sets according to Telenti et al. [19].

PCR reactions were conducted using 10 mM Tris-HCl (pH 8.3), 50 mM KCl, 2 mM MgCl2, 0.2 mM dNTP mix, 0.1 U μl-1 Taq polymerase, 0.5 μM of each of the primers, DNA template, and nuclease-free water. The PCR cycle conditions for amplifying the two genes (*IS6110* and *hsp65*) were as follows: 95°C for 4 min, followed by 30 cycles of 94°C for 30s, 65°C for 30s, and 72°C for 50s, with a final extension at 72°C for 10 min. To ensure the accuracy of the PCR, DNA from *M. tuberculosis H37Rv* and nuclease-free water (Sinaclon, BioScience) were utilized as positive and negative controls, respectively.

Analysis of 3μl of the PCR products (amplicon) was performed by electrophoresis on a 1.5% agarose gel. Following electrophoresis, the gel was stained with GelRed™ DNA stain, and the fragments were visualized under UV light using a gel documentation system (Gel Doc, ATP Co).

### 2.4. Sequencing analysis

Sequence analysis of the *hsp65* gene was utilized for the molecular identification of clinical isolates. PCR products were purified using a QIA quick PCR purification kit (QIAGEN, Germany) and then subjected to Sanger sequencing at Macrogen Corporation (Korea). The sequencing data for the *hsp65* gene were analyzed using DNASTAR Lasergene software (version 7.1).

The sequences were compared with similar sequences of the organisms in GenBank using the BLAST online software of the National Center for Biotechnology Information (https://blast.ncbi.nlm.nih.gov/Blast.cgi). NTM species identification was confirmed if a 97% match was achieved [20].

### 2.5. Phylogenetic analyses

Molecular phylogenetic analyses were conducted using MEGA XI (Molecular Evolutionary Genetics Analysis, version 11.0.13) [21].

### 2.6. Submission of nucleotide sequence data to GenBank

Nucleotide sequences were submitted to GenBank to obtain accession numbers. The submission of nucleotide sequences was conducted through the web-based BankIt platform (https://www.ncbi.nlm.nih.gov/WebSub/), and GenBank staff assigned accession numbers upon receipt. The GenBank accession numbers for the nucleotide sequences can be found in Table 3.

**Table 3.**
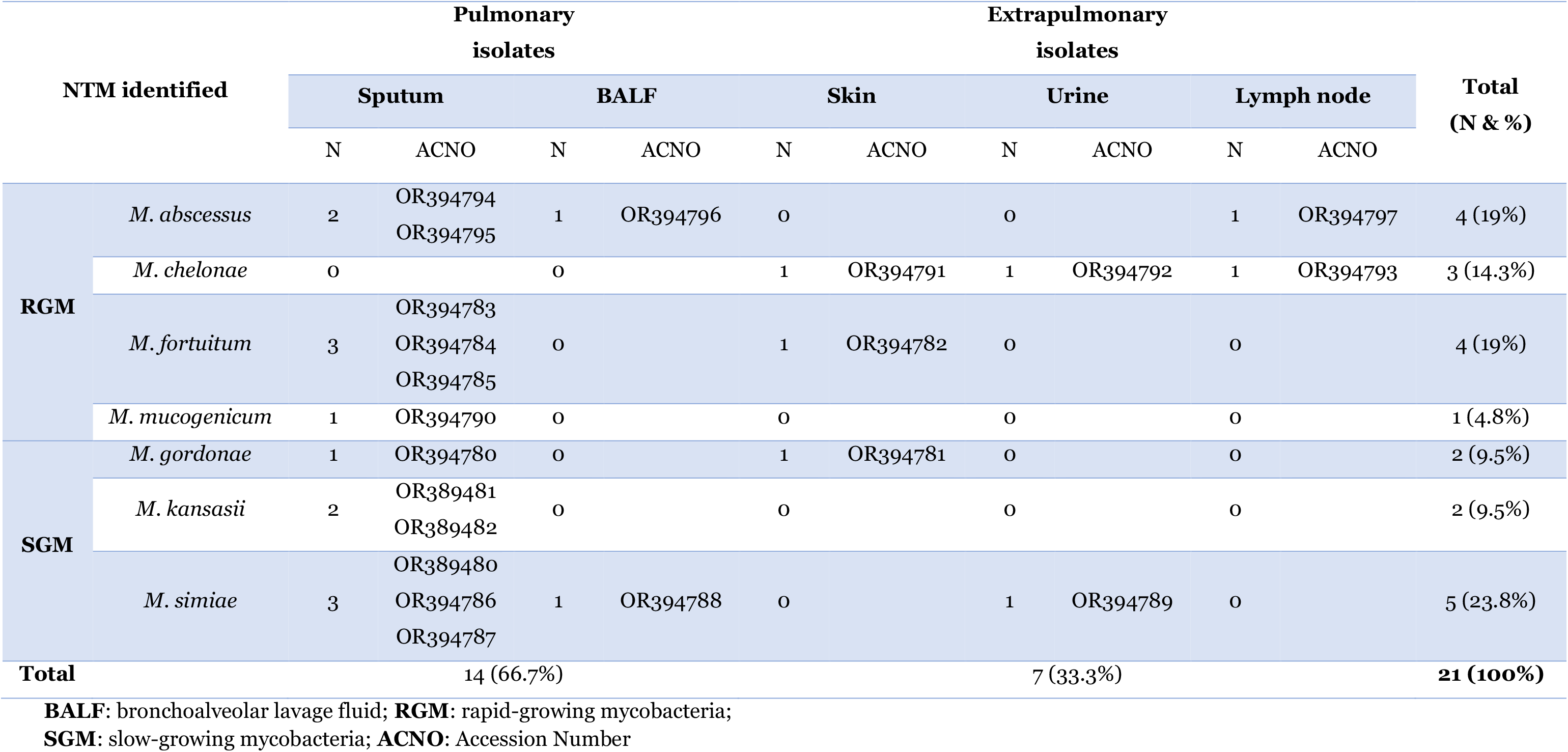
Distribution of NTM from clinical specimens with Accession numbers.

## 3. Results

Twenty-one NTM species were isolated from 230 clinical specimens (14 NTM from pulmonary specimens and 7 from extrapulmonary specimens) between 2021 and 2022. Of these, 12 (57.14%) were rapid-growing mycobacteria (RGM) and 9 (42.85%) were slow-growing mycobacteria (SGM). Overall, *M. simiae* (5/21, 24%) was the most frequently encountered species, followed by *M. abscessus* and *M. fortuitum*, each accounting for 4 out of the 21 isolates (19%). *M. chelonae* was present in 3 out of the 21 isolates (14.3%), while *M. kansasii* and *M. gordonae* were each identified in 2 out of the 21 isolates (9.5%), and *M. mucogenicum* was found in 1 out of the 21 isolates (4.7%). Among the 230 clinical specimens, 2 (0.87%) were confirmed as MTC using biochemical and molecular tests. Figure 1 displays the electrophoresis product of the 123 bp amplification of the *IS6110* sequence of MTC by PCR.

**Figure 1.**
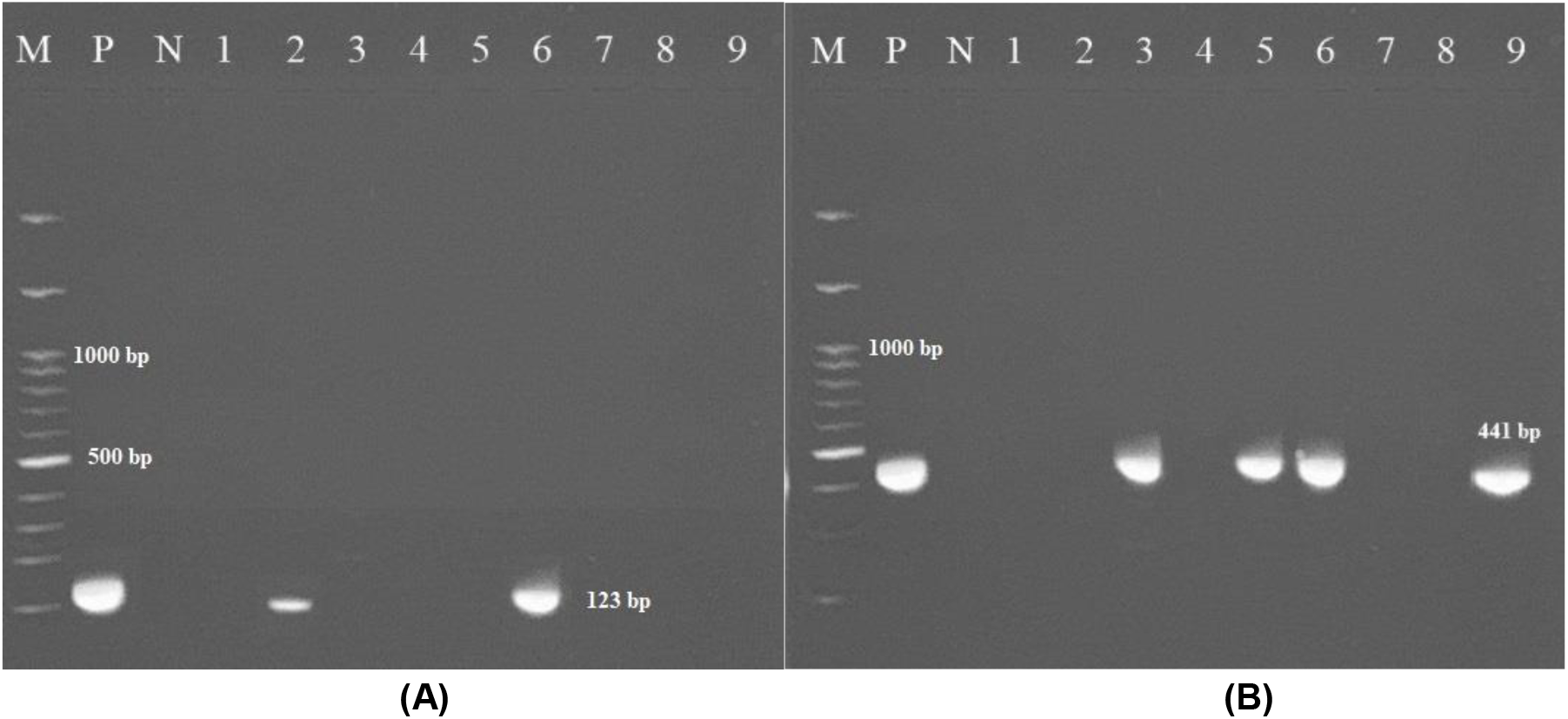
(A): Amplification of the 123 bp product of *M. tuberculosis* by PCR. PCR products (amplicon) were analyzed by electrophoresis on 1.5% gel agarose. (M: represents 100 bp DNA ladder; N: negative control; P: positive control DNA (*M. tuberculosis* strain *H37Rv*); 2, 6 clinically positive MTB specimens; 1, 3, 4, 5, 7, 8, 9 non-MTB specimens. (B): Figure to the right is 441-bp of the *hsp65* gene (M: Marker, P: positive, N: Negative); 3, 5, 6 clinically positive NTM specimens.

In the present study, we identified seven species of NTM in clinical specimens. *Mycobacterium avium* complex (MAC) was not detected in any of the specimens. Table 3 presents the specifics of pulmonary and extrapulmonary clinically significant NTM isolates and their distribution among patients based on specimen type. It is noteworthy that *M. mucogenicum* and *M. kansasii* were exclusively isolated from sputum specimens.

As can be seen in Table 3, 66.7% (14/21) of the identified NTM species were of pulmonary origin and 33.3% (7/21) were of extrapulmonary origin. Most species were isolated from sputum. Among pulmonary specimens, NTMs were identified in 12 (57.14%) sputum and two (9.5%) BALF. Among extrapulmonary specimens, NTMs were identified in three (14.3%) skin, two (9.5%) urine, and two (9.5%) lymph node samples.

The molecular phylogenetic and molecular evolutionary analysis of these species based on the *hsp65* gene sequences was conducted using MEGA XI version 11.0.13 software [21], and the representative result is shown in Figure 2. The *M. tuberculosis* strain *H37Rv* sequence was used as an out-group species. *M. kansasii, M. gordonae*, and *M. abscessus* strains showed high genetic diversity (Fig. 2). One strain of *M. abscessus* (obtained from sputum specimen No. 58) was genetically distinct from other identified *M. abscessus* species, as indicated by a star (*) in Figure 2.

**Figure 2.**
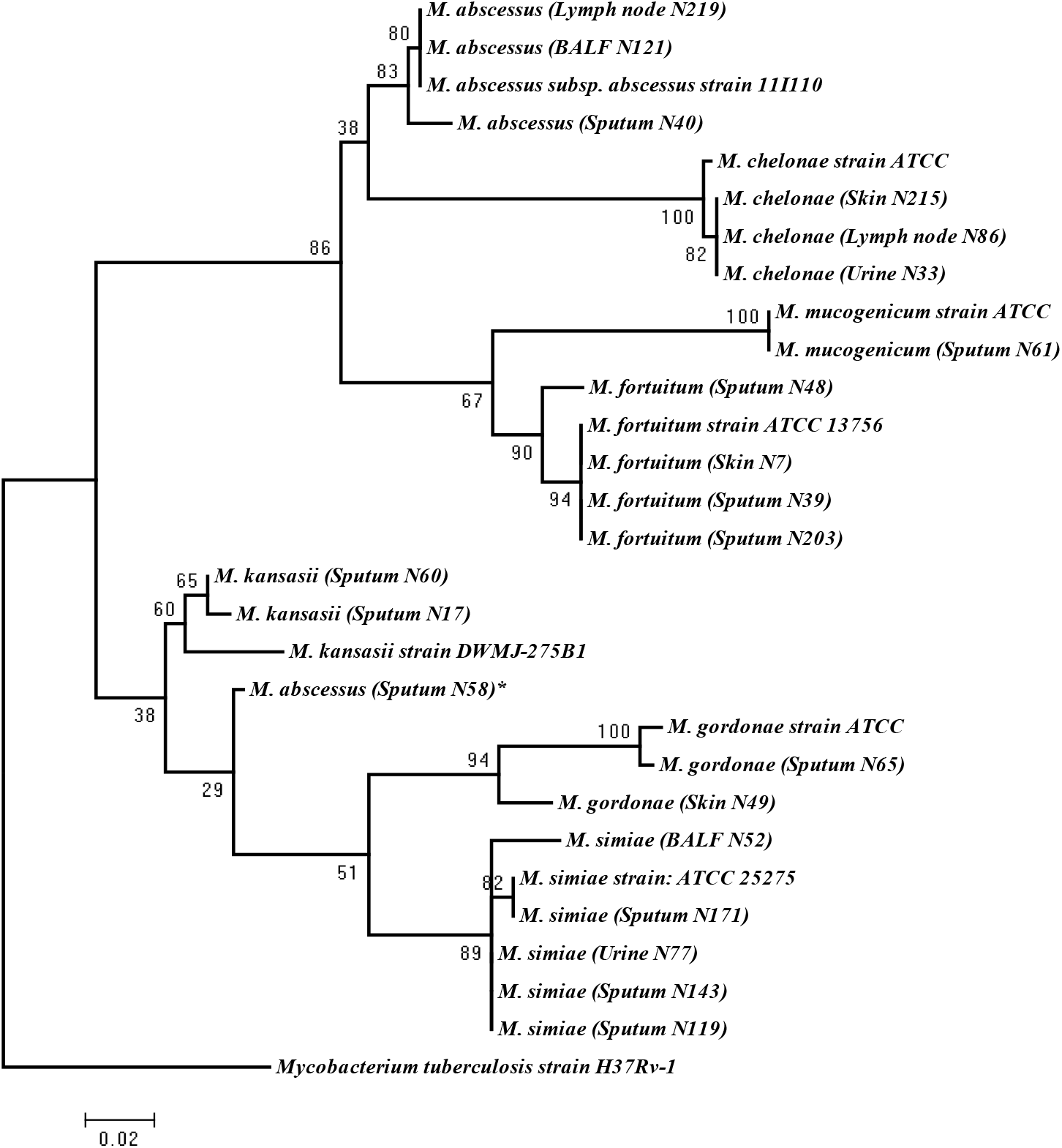
Phylogenetic tree based on the *hsp65* gene sequences of the isolates from the patients with suspected tuberculosis in East Azerbaijan province, northwest of Iran. A Maximum Likelihood tree was created, bootstrapped 100 times, and visualized with MEGA XI software (version 11.0.13) [21]. *M. tuberculosis* strain *H37Rv* was used as the out-group. The scale bar represents a 0.020 difference in nucleotide sequences.

## 4. Discussion

The increase in the outbreak of NTM infections in cases of suspected TB is a major global challenge [22]. NTM infection is typically believed to be a result of exposure to environmental sources such as water, soil, and dust. According to studies conducted in Iran, nontuberculous mycobacteria account for 5% to 10% of mycobacterial infections [23]. In this study, NTMs were identified in 9.13% (21/230) of patients with suspected tuberculosis, while only two MTB species were isolated from 230 clinical specimens. Therefore, it can be concluded that the frequency of infections caused by nontuberculous species is higher than that of tuberculosis. The prevalence of NTM in the study conducted by Hoza et al. in Tanzania was also found to be 9.7%, which is consistent with the results of the present study [24]. Furthermore, the prevalence of NTM in patients with suspected tuberculosis in the study conducted by Busatto et al. was 7% [22].

In this study, the diversity of NTM species was high, with 21 different species being isolated. Among these species, *M. simiae* was the most common, accounting for 23.8% of all isolates. *M. simiae* is a prevalent NTM species in Iran and has recently been recognized as an emerging pathogen [25,26,27]. It is notable that *M. simiae* is the only niacin-positive NTM that can be mistaken for MTB due to similar clinical symptoms [28]. These findings highlight the importance of investigating NTM species in patients suspected of having TB. Following *M. simiae, M. fortuitum*, and *M. abscessus* were the next most frequently isolated species in our study. Similar findings have been reported in other studies, such as those conducted by Sharma et al. [29], and Shenai et al.[30] in India, Albayrak et al. in Turkey [31], and Ahmed et al. in Pakistan [32]. In Taiwan, Greece, and the United Kingdom, *M. fortuitum* is the second most commonly isolated NTM species after members of the *M. avium* complex (MAC), with percentages of 23%, 21%, and 20%, respectively [33].

In contrast to other studies that have identified MAC as the dominant NTM species [34,35], our study did not isolate any MAC species from the specimens. Instead, we observed a prevalence of species such as *M. abscessus, M. chelonae, M. fortuitum, M. mucogenicum, M. gordonae, M. kansasii*, and *M. simiae* in East Azerbaijan, Iran.

Another study with similar findings was conducted by Gharbi et al. [36] in Tunisia. They investigated the presence of NTM in specimens of individuals suspected of having pulmonary tuberculosis in Northern Tunisia. In their study, they isolated nine species of NTM, including *M. kansasii, M. fortuitum, M. novocastrense, M. chelonae, M. gordonae, M. gadium, M. peregrinum, M. porcinum*, and *M. flavescens*. Like our study, they also did not isolate any MAC species from their specimens. In the Ide et al. study [14] in Nagasaki, Japan, *M. intracellulare* was the most common pathogen. In the Mwangi et al. study [20] in Kenya, the dominant NTM species was *Mycobacterium avium* complex. In the Mertaniasih et al. study [37], the most prevalent species were in the *M. avium-intracellulare* complex. In the Dadheech et al. study [38], *M. fortuitum* and *M. abscessus* were the most prevalent species. A study conducted across 30 predominantly European countries found that *M. xenopi* was most common in Hungary, *M. abscessus* complex (MABC) was most common in South Korea and Taiwan, and *M. kansasii* was most prevalent in Poland and Slovakia [39].

Therefore, it can be concluded that the distribution of NTM species varies in different geographical locations [39]. Our findings demonstrate a regional variation in the diversity of NTM species and suggest a potential influence of different geographical or environmental landscapes within East Azerbaijan province. NTMs frequently cause disease that is clinically indistinguishable from tuberculosis (TB). Therefore, it is crucial to accurately identify NTM species, understand their distribution patterns, and determine the predominant species in different geographical regions. This is important because treatment strategies for NTM-related diseases vary. The *hsp65* gene, present in all mycobacteria, has more polymorphism than the *16S rRNA* gene sequence and is useful for identifying genetically related species [18].

In this study, we used *hsp65* gene sequencing to identify nontuberculous mycobacteria species. This molecular technique has previously been successful in differentiating NTM species [6,18,40]. Our *hsp65* gene sequencing results confirmed that each species has a distinct and consistent sequence, as shown in Figure 3. The analysis presented in Figure 3 fully characterizes the nucleotide differences along the length of the *hsp65* gene. Interestingly, these differences are particularly abundant in two regions (97 to 106 and 178 to 205), indicating that these regions are the hypervariable regions of the *hsp65* gene.

**Figure 3.**
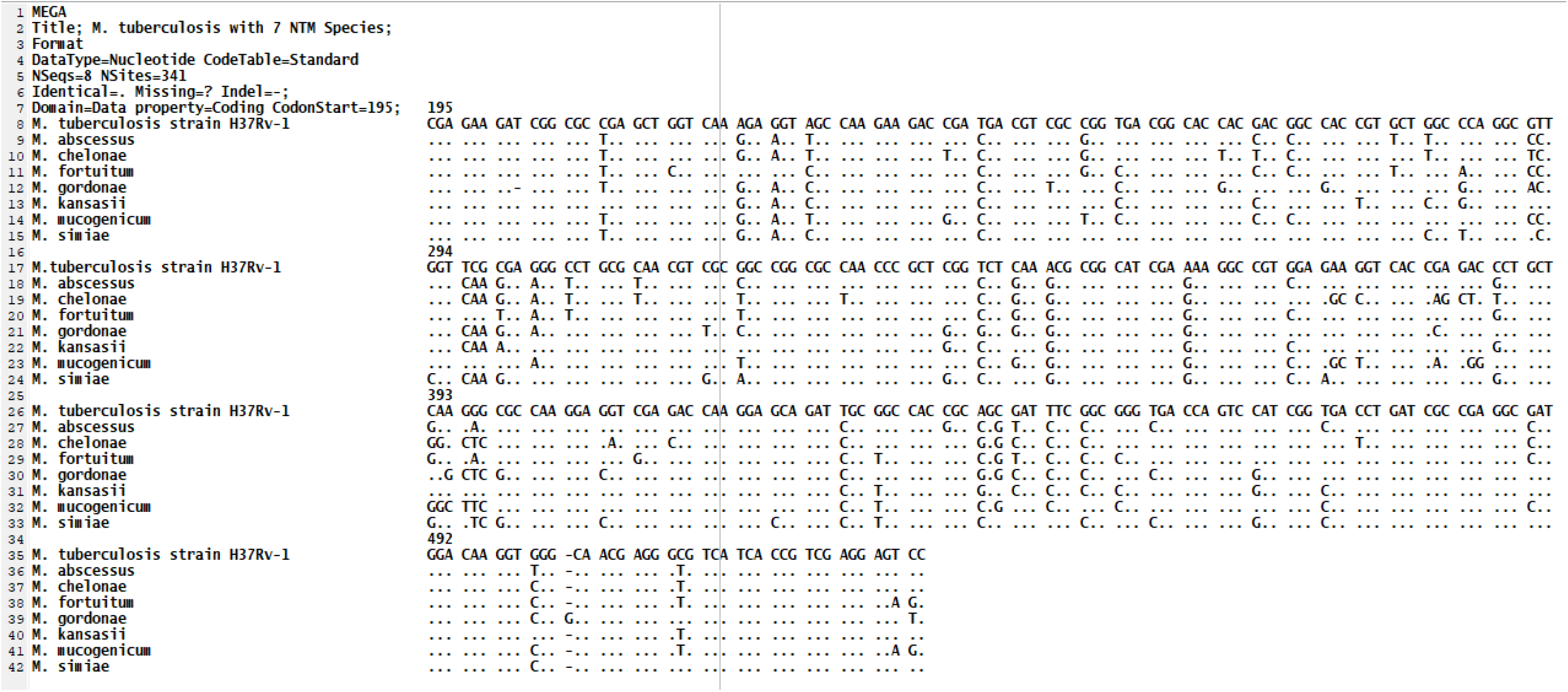
Alignment of partial sequence of the *hsp65* gene from seven different species. *M. tuberculosis* strain *H37Rv1* was used as differing from those of the MTB sequence are indicated; dots indicate identity. The first nucleotide shown corresponds to position 195 of the published sequence from MTB. an out-group; nucleotides

Among rapid-growing mycobacteria, *M. mucogenicum* and *M. fortuitum* showed a high degree of similarity. On the other hand, *M. chelonae* and *M. mucogenicum* were the two species from RGM with the highest degree of difference. For the similarity and difference grades of other species, please refer to Figure 4.

**Figure 4.**
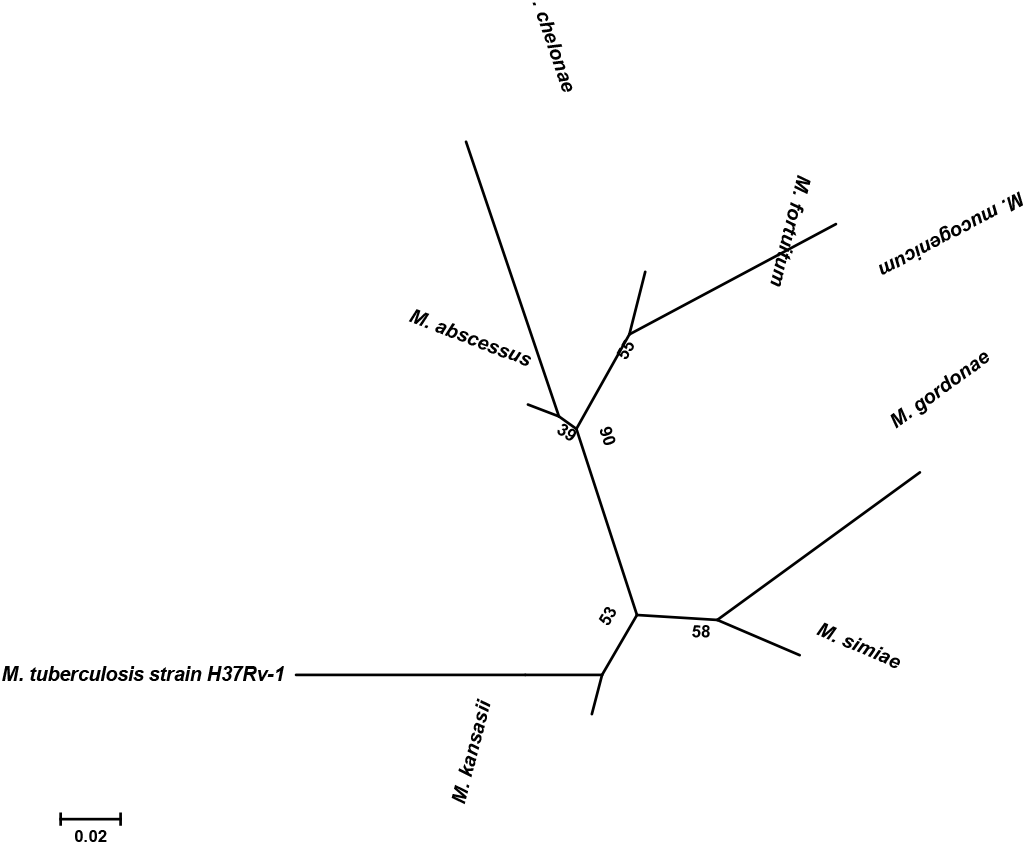
Molecular Phylogenetic analysis by Maximum Likelihood method based on *hsp65* sequences. The bar indicates a 0.02 estimated sequence divergence.

## 5. Conclusion

This study is the first to characterize the diversity of NTM in patients with suspected tuberculosis in east Azerbaijan province. It identified 21 different NTM species, with *M. simiae* being the most prevalent, followed by *M. fortuitum* and *M. abscessus*. Additionally, the study revealed a higher frequency of infections caused by nontuberculous species compared to tuberculosis. These findings emphasize the importance of investigating NTM species in suspected TB patients. In endemic areas of TB, NTM-PD can be misdiagnosed as pulmonary tuberculosis (TB), leading to repeated failures in anti-TB treatment and increased complications. This highlights the need for better diagnosis.

## Ethical Approval

Our study did not involve human participants, so ethical approval was not necessary. We only worked on bacteria.

## References

1. Zulu M, Monde N, Nkhoma P, et al. (2021) Nontuberculous Mycobacteria in Humans, Animals, and Water in Zambia: A Systematic Review. Frontiers in Tropical Diseases 2.

2. Salzer HJF, Chitechi B, Hillemann D, et al. (2020) Nontuberculous Mycobacterial Pulmonary Disease from Mycobacterium hassiacum, Austria. Emerg Infect Dis 26: 2776–2778.

3. Roshdi Maleki M, Moaddab SR, Samadi kafil H (2019) Hemodialysis waters as a source of potentially pathogenic mycobacteria (PPM). DESALINATION AND WATER TREATMENT.

4. Yu P, Yaoju T, Jin C, et al. (2017) Diversity of nontuberculous mycobacteria in eastern and southern China: a cross-sectional study. European Respiratory Journal 49: 1601429.

5. Desai AN, Hurtado R (2021) Nontuberculous Mycobacterial Infections. JAMA 325: 1574–1574.

6. Maleki MR, Kafil HS, Harzandi N, et al. (2017) Identification of nontuberculous mycobacteria isolated from hospital water by sequence analysis of the hsp65 and 16S rRNA genes. J Water Health 15: 766–774.

7. Cowman S, van Ingen J, Griffith DE, et al. (2019) Non-tuberculous mycobacterial pulmonary disease. European Respiratory Journal 54: 1900250.

8. Cook JL (2010) Nontuberculous mycobacteria: opportunistic environmental pathogens for predisposed hosts. Br Med Bull 96: 45–59.

9. Porvaznik I, Solovič I, Mokrý J (2017) Non-Tuberculous Mycobacteria: Classification, Diagnostics, and Therapy. In: Pokorski M, editor. Respiratory Treatment and Prevention. Cham: Springer International Publishing. pp. 19–25.

10. Adjemian J, Olivier KN, Seitz AE, et al. (2012) Prevalence of nontuberculous mycobacterial lung disease in U.S. Medicare beneficiaries. American journal of respiratory and critical care medicine 185: 881–886.

11. Morimoto K, Iwai K, Uchimura K, et al. (2014) A steady increase in nontuberculous mycobacteriosis mortality and estimated prevalence in Japan. Ann Am Thorac Soc 11: 1–8.

12. Shah NM, Davidson JA, Anderson LF, et al. (2016) Pulmonary Mycobacterium avium-intracellulare is the main driver of the rise in non-tuberculous mycobacteria incidence in England, Wales and Northern Ireland, 2007-2012. BMC Infect Dis 16: 195.

13. Thomson R, Donnan E, Konstantinos A (2017) Notification of Nontuberculous Mycobacteria: An Australian Perspective. Ann Am Thorac Soc 14: 318–323.

14. Ide S, Nakamura S, Yamamoto Y, et al. (2015) Epidemiology and clinical features of pulmonary nontuberculous mycobacteriosis in Nagasaki, Japan. PLoS One 10: e0128304.

15. Bradner L, Robbe-Austerman S, Beitz DC, et al. (2013) Chemical decontamination with N-acetyl-L-cysteine-sodium hydroxide improves recovery of viable Mycobacterium avium subsp. paratuberculosis organisms from cultured milk. J Clin Microbiol 51: 2139–2146.

16. van Soolingen D, Hermans PW, de Haas PE, et al. (1991) Occurrence and stability of insertion sequences in Mycobacterium tuberculosis complex strains: evaluation of an insertion sequence-dependent DNA polymorphism as a tool in the epidemiology of tuberculosis. J Clin Microbiol 29: 2578–2586.

17. Kabir S, Uddin MKM, Chisti MJ, et al. (2018) Role of PCR method using IS6110 primer in detecting Mycobacterium tuberculosis among the clinically diagnosed childhood tuberculosis patients at an urban hospital in Dhaka, Bangladesh. Int J Infect Dis 68: 108–114.

18. Kim H, Kim SH, Shim TS, et al. (2005) Differentiation of Mycobacterium species by analysis of the heat-shock protein 65 gene (hsp65). Int J Syst Evol Microbiol 55: 1649–1656.

19. Telenti A, Marchesi F, Balz M, et al. (1993) Rapid identification of mycobacteria to the species level by polymerase chain reaction and restriction enzyme analysis. J Clin Microbiol 31: 175–178.

20. Mwangi ZM, Mukiri NN, Onyambu FG, et al. (2022) Genetic Diversity of Nontuberculous Mycobacteria among Symptomatic Tuberculosis Negative Patients in Kenya. The International Journal of Mycobacteriology 11: 60–69.

21. Tamura K, Stecher G, Kumar S (2021) MEGA11: Molecular Evolutionary Genetics Analysis Version 11. Molecular Biology and Evolution 38: 3022–3027.

22. Busatto C, Vianna JS, Silva ABS, et al. (2020) Nontuberculous mycobacteria in patients with suspected tuberculosis and the genetic diversity of Mycobacterium avium in the extreme south of Brazil. J Bras Pneumol 46: e20190184.

23. Nasiri MJ, Dabiri H, Darban-Sarokhalil D, et al. (2015) Prevalence of Non-Tuberculosis Mycobacterial Infections among Tuberculosis Suspects in Iran: Systematic Review and Meta-Analysis. PLoS One 10: e0129073.

24. Hoza AS, Mfinanga SG, Rodloff AC, et al. (2016) Increased isolation of nontuberculous mycobacteria among TB suspects in Northeastern, Tanzania: public health and diagnostic implications for control programmes. BMC Res Notes 9: 109.

25. Hashemi-Shahraki A, Darban-Sarokhalil D, Heidarieh P, et al. (2013) Mycobacterium simiae: a possible emerging pathogen in Iran. Jpn J Infect Dis 66: 475–479.

26. Heidarieh P, Mirsaeidi M, Hashemzadeh M, et al. (2016) In Vitro Antimicrobial Susceptibility of Nontuberculous Mycobacteria in Iran. Microb Drug Resist 22: 172–178.

27. Maoz C, Shitrit D, Samra Z, et al. (2008) Pulmonary Mycobacterium simiae infection: comparison with pulmonary tuberculosis. Eur J Clin Microbiol Infect Dis 27: 945–950.

28. Bhalla GS, Sarao MS, Kalra D, et al. (2018) Methods of phenotypic identification of non-tuberculous mycobacteria. Practical Laboratory Medicine 12: e00107.

29. Sharma SK, Upadhyay V (2020) Epidemiology, diagnosis & treatment of non-tuberculous mycobacterial diseases. Indian J Med Res 152: 185–226.

30. Shenai S, Rodrigues C, Mehta A (2010) Time to identify and define non-tuberculous mycobacteria in a tuberculosis-endemic region. Int J Tuberc Lung Dis 14: 1001–1008.

31. Albayrak N, Simşek H, Sezen F, et al. (2012) [Evaluation of the distribution of non-tuberculous mycobacteria strains isolated in National Tuberculosis Reference Laboratory in 2009-2010, Turkey]. Mikrobiyol Bul 46: 560–567.

32. Ahmed I, Jabeen K, Hasan R (2013) Identification of non-tuberculous mycobacteria isolated from clinical specimens at a tertiary care hospital: a cross-sectional study. BMC Infect Dis 13: 493.

33. Hoefsloot W, van Ingen J, Andrejak C, et al. (2013) The geographic diversity of nontuberculous mycobacteria isolated from pulmonary samples: an NTM-NET collaborative study. Eur Respir J 42: 1604–1613.

34. Clarke T, Brinkac L, Manoranjan J, et al. (2020) Typing and classification of non-tuberculous mycobacteria isolates [version 1; peer review: 1 approved with reservations]. City.

35. Chalmers JD, Balavoine C, Castellotti PF, et al. (2020) European Respiratory Society International Congress, Madrid, 2019: nontuberculous mycobacterial pulmonary disease highlights. ERJ Open Research 6: 00317–02020.

36. Gharbi R, Mhenni B, Ben Fraj S, et al. (2019) Nontuberculous mycobacteria isolated from specimens of pulmonary tuberculosis suspects, Northern Tunisia: 2002-2016. BMC infectious diseases 19: 819–819.

37. Mertaniasih NM, Kusumaningrum D, Koendhori EB, et al. (2017) Nontuberculous mycobacterial species and Mycobacterium tuberculosis complex coinfection in patients with pulmonary tuberculosis in Dr. Soetomo Hospital, Surabaya, Indonesia. Int J Mycobacteriol 6: 9–13.

38. Dadheech M, Malhotra AG, Patel S, et al. (2023) Molecular Identification of Non-tuberculous Mycobacteria in Suspected Tuberculosis Cases in Central India. Cureus 15.

39. Hoefsloot W, Van Ingen J, Andrejak C, et al. (2013) The geographic diversity of nontuberculous mycobacteria isolated from pulmonary samples: an NTM-NET collaborative study. European Respiratory Journal 42: 1604–1613.

40. Escobar-Escamilla N, Ramírez-González JE, González-Villa M, et al. (2014) hsp65 Phylogenetic Assay for Molecular Diagnosis of Nontuberculous Mycobacteria Isolated in Mexico. Archives of Medical Research 45: 90–97.

